# The use of CRISPR-Cas Selective Amplicon Sequencing (CCSAS) to reveal the eukaryotic microbiome of metazoans

**DOI:** 10.1101/2020.06.02.130807

**Authors:** Kevin Xu Zhong, Anna Cho, Christophe M. Deeg, Amy M. Chan, Curtis A. Suttle

**Author notes:** Correspondence to: Curtis Suttle, Department of Earth, Ocean & Atmospheric Sciences, University of British Columbia, #2020-2207 Main Mall, Vancouver, BC, Canada V6T 1Z4, Kevin Xu Zhong, Department of Earth, Ocean & Atmospheric Sciences, University of British Columbia, #2020-2207 Main Mall, Vancouver, BC, Canada V6T 1Z4.

## Abstract

Characterization of the eukaryotic microbiome is required to understand the role of microbial communities in health and disease. Such investigation relies on sequencing 18S ribosomal RNA genes (rDNA), which serve as taxonomic markers; however, this is compromised by contaminating host rDNA sequences. To overcome this problem, we developed CRISPR-Cas Selective Amplicon Sequencing (CCSAS), a high-resolution and efficient approach for characterizing eukaryotic microbiomes. CCSAS uses taxon-specific single-guide RNA (sgRNA) to direct Cas9 to cut 18S rDNA sequences of the host. Validation shows that >96.5% of rDNA amplicons from ten model organisms were cleaved, while rDNA from protists and fungi were unaffected. In oyster spat, CCSAS resolved ∼8.5-fold more taxa, and several additional major phylogenetic groups when compared to the best available alternative approach. We designed taxon-specific sgRNA for ∼16,000 metazoan and plant taxa, making CCSAS widely available for characterizing eukaryotic microbiomes that have largely been neglected because of methodological challenges.

## Introduction

There is an explosion in interest and understanding of how the composition of the microbiome affects the health of plants^1-2^ and animals^3-6^, including humans^7-9^. For example, the composition of the microbiome is associated with both positive and adverse health effects in humans^10-14^. As well, a wide span of biological, ecological and evolutionary questions have been addressed through microbiome studies^6-8, 15-17^.

Deep sequencing of ribosomal RNA gene (rDNA) fragments is widely used to characterize the microbiome associated with a wide range of organisms^18-21^. Yet, to date, our knowledge on the eukaryotic component of the microbiome, particularly protists, is relatively limited compared to that of prokaryotes^6, 21-25^. This is due to the challenge in profiling host-associated eukaryotic microbes, as the standard “universal” primers^26^ used to amplify 18S rDNA sequences from eukaryotic microbes also amplify the host 18S rDNA, which will predominate the sequencing library^18^. Sampling strategies to minimize host-tissue contamination, including using fecal samples for the gut microbiome, or using gentle brushing to sample the skin microbiome are not possible when the host organism is too small or the microbes are too closely associated with host tissue. Other strategies such as using taxon-specific primers to target microbial 18S sequences (e.g. non-metazoan primers^27^), using other marker genes such as the ITS region of fungi^28^, or using blocking primers^29^ to prevent the amplification of host 18S rDNA sequences have significant challenges when trying to target the entire eukaryotic microbial community. For example, blocking primers^29^ focus on a short fragment within the hypervariable region that has a reduced taxonomic resolution compared to “universal” 18S primers. Second, designing a primer that blocks amplification of host DNA, but discriminates a wide variety of taxa is often not possible and hinders the broad application of this approach. Third, even with blocking primers, a large proportion of sequences can be host-derived (e.g. up to 92% in coral, 42% in krill, and 45% in fish) as briefly reviewed^30^. Consequently, studies on the eukaryotic microbiome are languishing in the absence of an approach that allows effective amplification of microeukaryotic 18S sequences with minimal contamination from host sequences.

Clustered regularly interspaced short palindromic repeats (CRISPR) and the CRISPR-associated protein 9 (Cas9) system offers bacteria and archaea adaptive immunity against viruses and plasmids by cleaving invading double-stranded (ds) DNA^31^. The sequence-specific cleavage is performed by Cas9 endonuclease in the presence of guide RNA (gRNA). This gRNA is a duplex comprising a trans-activating RNA (tracrRNA) that is a scaffold for binding the Cas9 protein, and an approximately 20 nucleotide (nt) crispr RNA (crRNA) guide sequence that is complementary to the DNA target site^32-35^. Cas9 can be programmed to target any DNA sequence by modifying the 20-nt guide sequence^35-36^. Due to its precision in DNA cutting, the simplicity in programming and the ability to artificially fuse the gRNA duplex (tracrRNA-crRNA) into a single-guide RNA (sgRNA)^35^, CRISPR-Cas9 has emerged as a powerful biotechnological tool in a wide variety of applications^36-37^. We used this approach and designed a custom sgRNA to direct Cas9 to specifically cut host 18S rDNA sequences in the region targeted by “universal” primers. The cleaved host 18S fragments contain only a 3’ or 5’ primer binding region, resulting in short single-stranded (ss) DNA products produced by PCR, which are then removed by size-selection using SPRI magnetic beads during the preparation of the sequencing library. This results in a library highly enriched in 18S amplicons from protists and fungi, allowing for high-resolution surveys of the taxonomic composition of the eukaryotic microbes associated with any eukaryotic host. This sequencing library preparation method is termed as CRISPR-Cas Selective Amplicon Sequencing (CCSAS).

## Results

### Design of the taxon-specific sgRNA

The key to CCSAS is the 20-nt guide sequence of gRNA that directs Cas9 to cut the 18S rDNA sequences of the host, but not those of the associated microeukaryotes. We developed CasOligo (https://github.com/kevinzhongxu/CasOligo), an R package that contains the algorithm Cas9.gRNA.oligo1(), which identifies 20-nt sequences in the 18S rDNA region spanned by “universal” primers that can serve as target-sites for gRNA, and which are compatible with the CRISPR-Cas9 system and specific to the 18S rRNA gene of the host but not to microeukaryotes (Supplementary Materials and Methods). This 20-nt gRNA-target-site oligonucleotide sequence, complementary to the sgRNA’s guide sequence, determines the specificity of the sgRNA and thereby the CRISPR-Cas action, and can be used to design and synthesize a taxon-specific sgRNA in the laboratory (Supplementary Materials and Methods).

To validate the design of sgRNA in taxon-specific cleavage, taxon-specific sgRNAs were designed and tested for each of the following ten model organisms: human (*Homo sapiens*), salmon *(Salmo salar*), shrimp (*Solenocera crassicornis*), chicken (*Gallus gallus domesticus*), cow (*Bos taurus*), mouse (*Mus musculus*), fruit fly (*Drosophila melanogaster*), rock cress (*Arabidopsis thaliana*), oyster (*Crassostrea gigas*) and the nematode (*Caenorhabditis elegans*), as well as being tested against a mock community composed of nine pathogenic protists and fungi (Supplementary Table 1 and 2). The results showed that the CRISPR-Cas9 treatment effectively cleaved the host 18S amplicons, while amplicons from the mock community of protists and fungi remained intact (Fig. 1). Comparisons using qPCR with and without CRISPR-Cas9 treatment showed that only 0.6% to 3.5% of the intact 18S amplicons remained after CRISPR-Cas9 cutting (Fig. 1D). Thus, the sgRNAs effectively targeted host sequences, while leaving sequences from microeukaryotes intact.

**Fig. 1.**
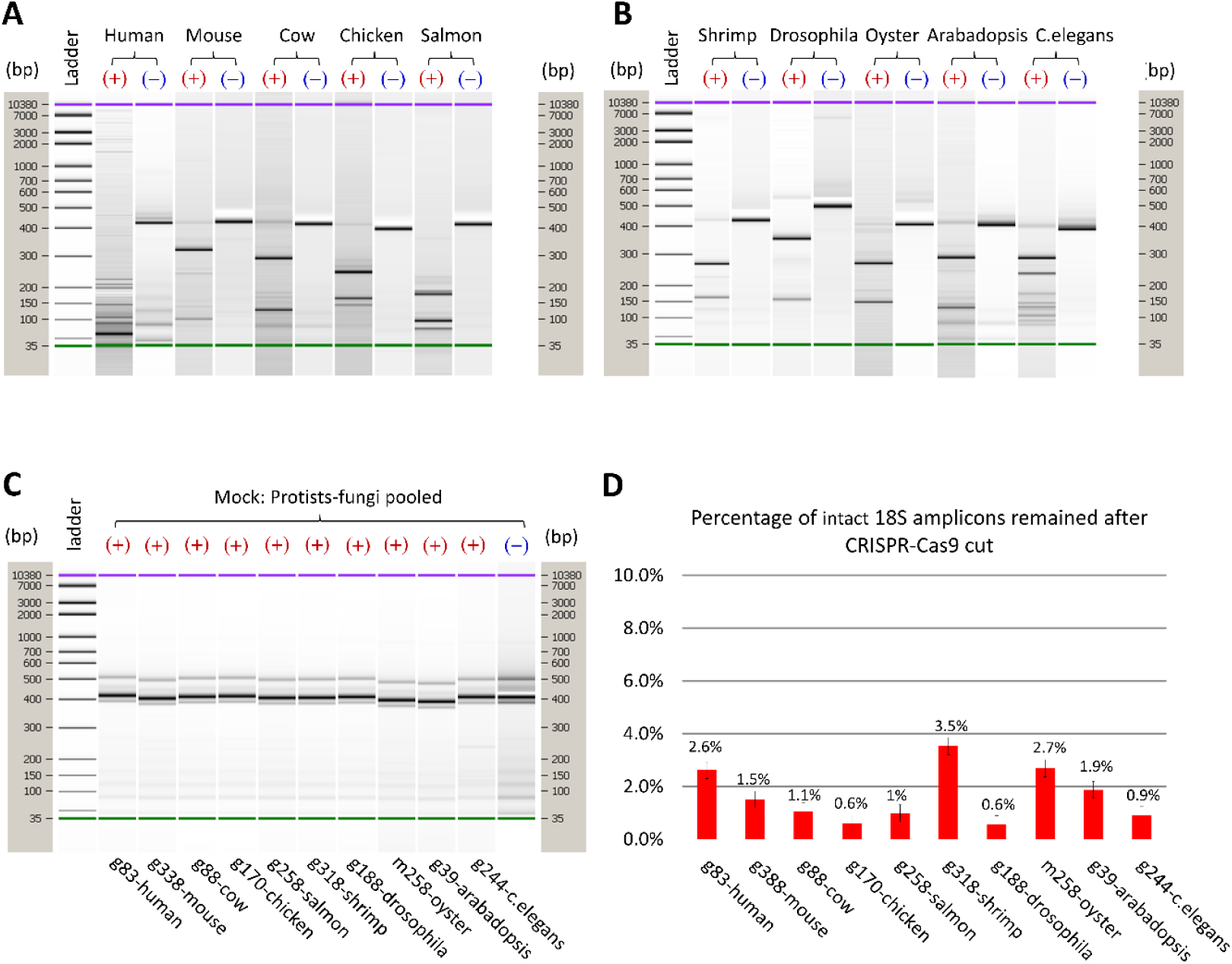
Gel images of 18S amplicons from ten model organisms (**A** and **B**) and a mock community of protists and fungi (**C**) to which Cas9 with the taxon-specific sgRNA (as shown in Supplementary Table 2) was either added (**+**) or not (**-**). Gel bands show the amplicon length in base pairs (bp) relative to a DNA ladder. Quantitative PCR was used to assess the remaining intact 18S amplicons from the model organisms after cutting with CRISPR-Cas9 (**D)**. The labels on the X-axes of C and D indicate the ID of the taxon-specific gRNAs and its corresponding host.

### Using CCSAS to reveal host-associated microeukaryotic populations

Cas9 in association with the host-specific sgRNA will specifically cut the amplified host 18S sequences, while leaving intact the DNA amplified from microeukaryotes. This allows for high-resolution characterization of the taxonomic composition of the microeukaryotic community with a fraction of the sequencing effort typically used. However, after Cas9 treatment 0.6% to 3.5% of the 18S amplicons remaining were still primarily host-derived, and these could still dominate the sequencing library (data not shown). Hence, to further reduce host-derived 18S rRNA gene sequences we introduced a two-step CCSAS approach (Fig. 2) which employs a two-step CRISPR-Cas9 procedure. First, Cas9 with a taxon-specific sgRNA that is complementary to the host 18S rRNA gene sequence at the 20-nt target-site is used to cut the host genomic 18S rDNA, and then the remaining uncut 18S sequences are amplified using PCR. Any amplification of the cut fragments yields short pieces of ssDNA that are removed during size-selection using SPRI magnetic beads (Fig. 2). Second, following the first size selection, another Cas9 cut, PCR amplification and size selection were conducted, resulting in almost the complete removal of host 18S amplicons, while leaving the protistan and fungal amplicons intact and enriched for sequencing (Fig. 2). This two-step CCSAS was used to examine the eukaryotic microbiome from eight different samples of oyster spat (*C. gigas*) collected from a hatchery that was experiencing mortality events.

**Fig. 2.**
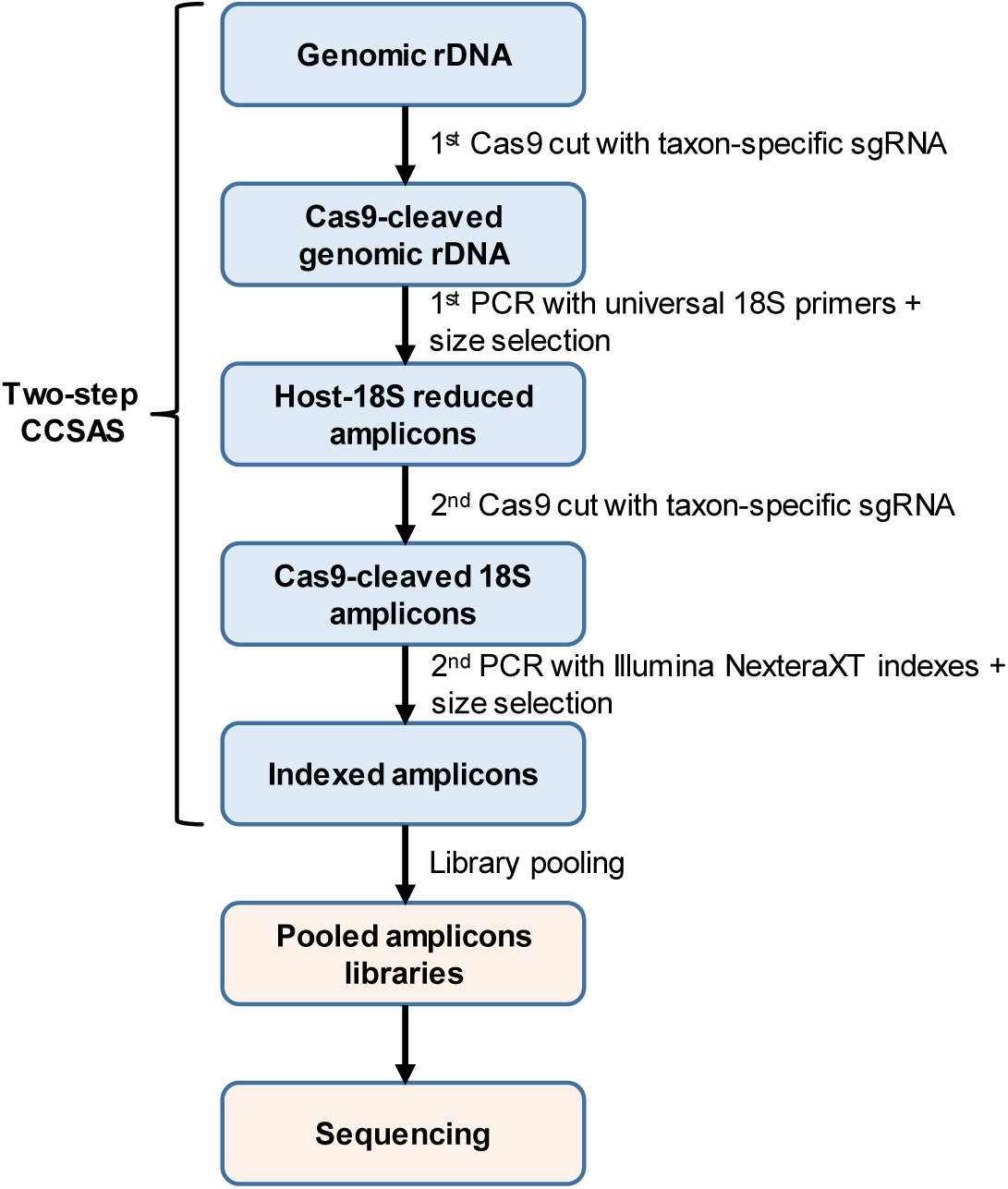
Workflow of the two-step CRISPR-Cas Selective Amplicon Sequencing (CCSAS) to study the host-associated eukaryotic microbiome.

The results showed almost the complete removal of oyster 18S amplicons by CCSAS, while leaving the protistan and fungal amplicons intact and highly enriched for sequencing (Fig. 3). Furthermore, the taxonomic resolution of the eukaryotic microbial community associated with the oysters was much higher when using CCSAS in conjunction with “universal” 18S primers than with the best available alternative approach using primers that target non-metazoan eukaryotes (Fig. 3, Fig. 4). Relative to using non-metazoan primers, CCSAS revealed ∼8.5-fold more unique taxa of microeukaryotes (Fig. 4B). Of these, ∼5.7-fold more were assigned to established taxa (Fig. 4C), such as Picozoa, Apusomonadidae, Chloroplastida, Centrohelida and Amoebozoa, as well as several groups within the Alveolata including Protalveolata, Dinoflagellata and Apicomplexa (Fig. 4D). Thus, using amplification with “universal” 18S primers combined with CCSAS provided much better taxonomic resolution of the eukaryotic microbiome, and more efficient sequencing compared to using non-metazoan primers. Moreover, with non-metazoan primers, most sequences were from metazoa (Fig. 4A), although the taxonomic resolution was not adequate to assign them to be from oysters (Fig. 3). In contrast, for CCSAS, the highest percentage of sequences from metazoa (mostly assigned to oysters and nematodes of order Monhysterida; Fig. 3) was only 7.4%, while in three out of eight samples, sequences from metazoa were undetectable (Fig. 4A).

**Fig. 3.**
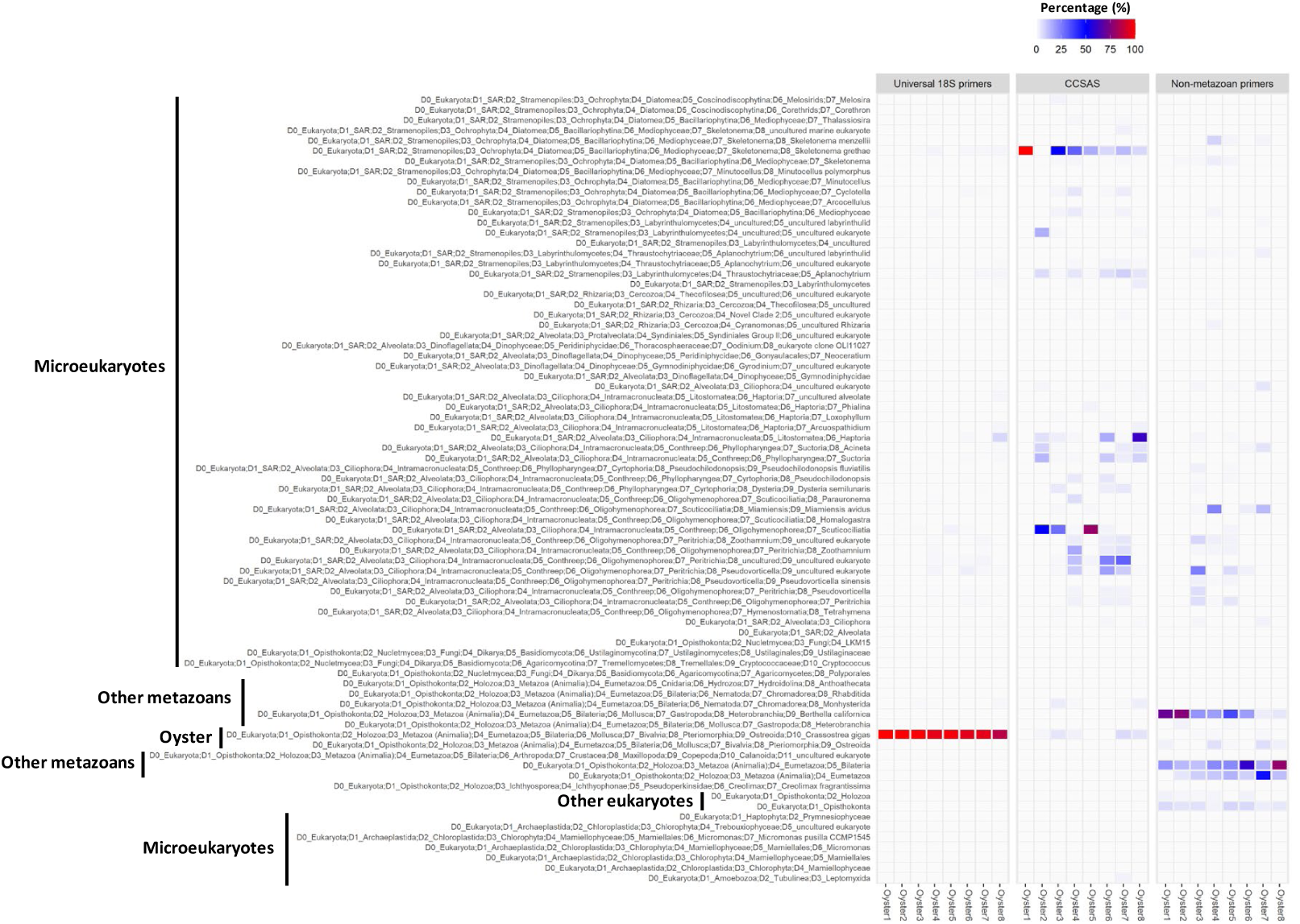
Eukaryotic taxa representing >1% of the sequences revealed by amplicon deep-sequencing of the 18S rRNA genes for oyster spat samples, using “**universal” 18S primers**^26^ (Supplementary Table 3), **non-metazoan primers**^27^ (Supplementary Table 3), as well as CRISPR-Cas Selective Amplicons Sequencing (**CCSAS**), which combined “universal” 18S primers and CRISPR-Cas9 with pacific-oyster-specific sgRNA m258 (Supplementary Table 2).

**Fig. 4.**
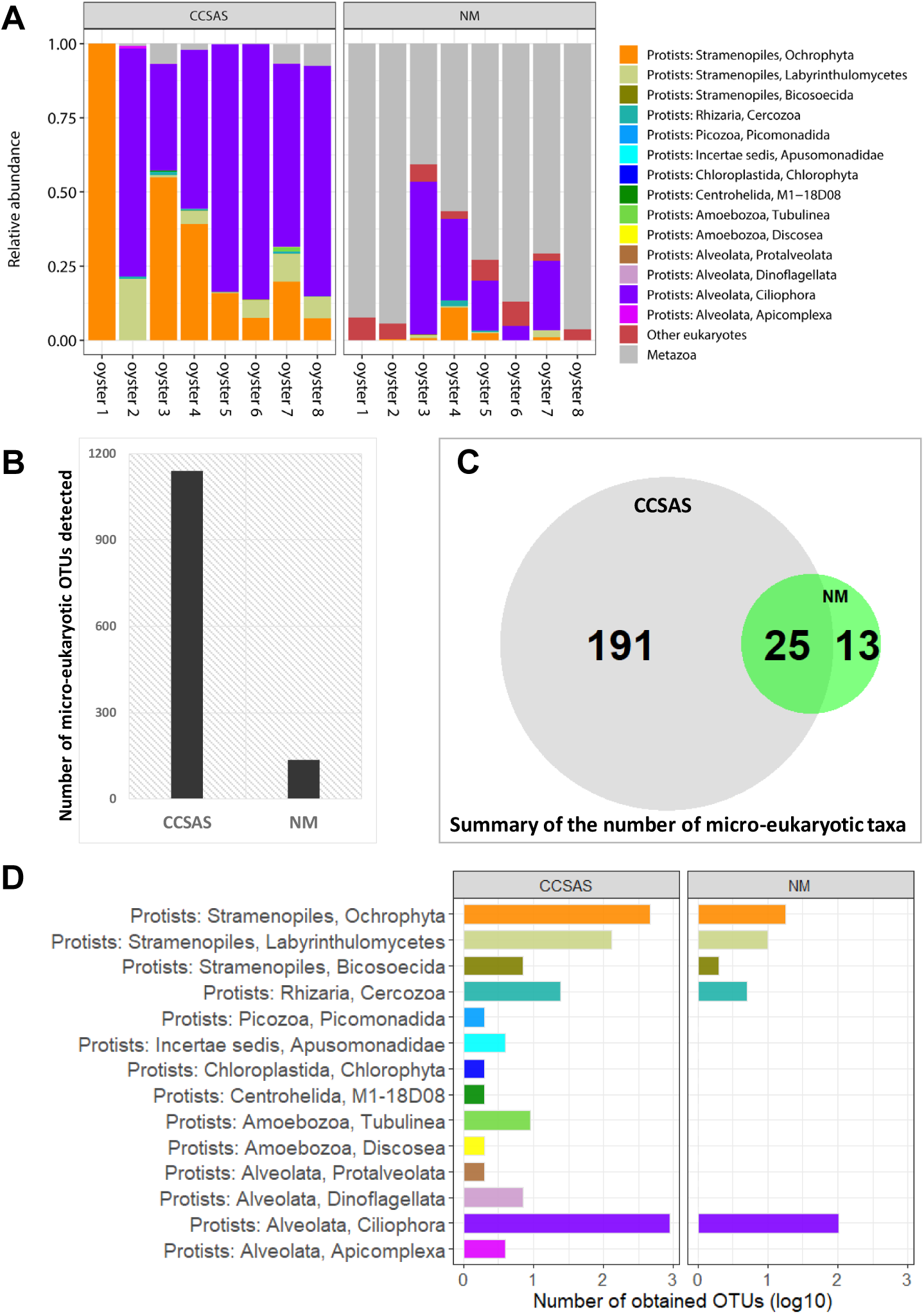
Distribution of eukaryotic groups (**panel A**) in eight oyster samples revealed using deep-sequencing of 18S amplicons of rRNA genes using non-metazoan primers (**NM**), and CCSAS that combines 18S “universal” primers and cleavage using CRISPR-Cas9 (**CCSAS**). The number of unique operational taxonomic units (OTUs) of microeukaryotes (**panel B**) and their assigned taxa (**panel C**) are summarized for the NM and CCSAS methods. **Panel D** compares the number of OTUs detected for different microeukaryotic groups using the CCSAS and NM methods.

### Database of gRNA-target-sites for metazoa and plants

To ease the application of CCSAS to the study of eukaryotic microbiomes in a wide range of metazoa and plants, we used CasOligo to identify the gRNA-target-sites for almost all 15907 metazoa and plant taxa (metaphyta of Embryophyta group) in the SILVA SSU database^38^ (*version v119, released on 24 July 2014*) (Fig. 5). This created a database of gRNA-target-sites, comprising, on average, 33 taxon-specific gRNA-target-sites per taxon (Supplementary Fig. 1), which provides a wide choice for desiging taxon-specific sgRNA for various host organisms. For each metazoan and plant species in SILVA, we revealed between 3 and 217 different gRNA-target-sites that are compatible with the CRISPR-Cas9 system (Supplementary Fig. 1). For 99.6% of the 15907 metazoan and plant species in SILVA, we found between 1 and 214 gRNA-target-sites that specifically target the 18S sequence for each taxon, but not protistan or fungal 18S sequences. Although it is not possible to design a “universal” sgRNA that targets all metazoa and plants but not microeukaryotes, some sgRNAs target broad taxonomic groups (Supplementary Fig. 2). For example, based on in-silico analysis, sgRNA_058534 targets 3099 species from 22 classes and families of Animalia, primarily 72.7% of the 4014 Insecta species in SILVA (Supplementary Fig. 2). CasOligo, also allows retrieval of the gRNA-target-site oligonucleotide sequence for specific taxa by entering the scientific name of the host species using the search.db.byname() function. Nonetheless, it is best to identify the taxon-specific gRNA-target-site based on the 18S rRNA gene sequence of the host, because the action of CRISPR-Cas9 is highly sequences-specific and the database does not cover all sequence variants for specific taxa.

**Fig. 5.**
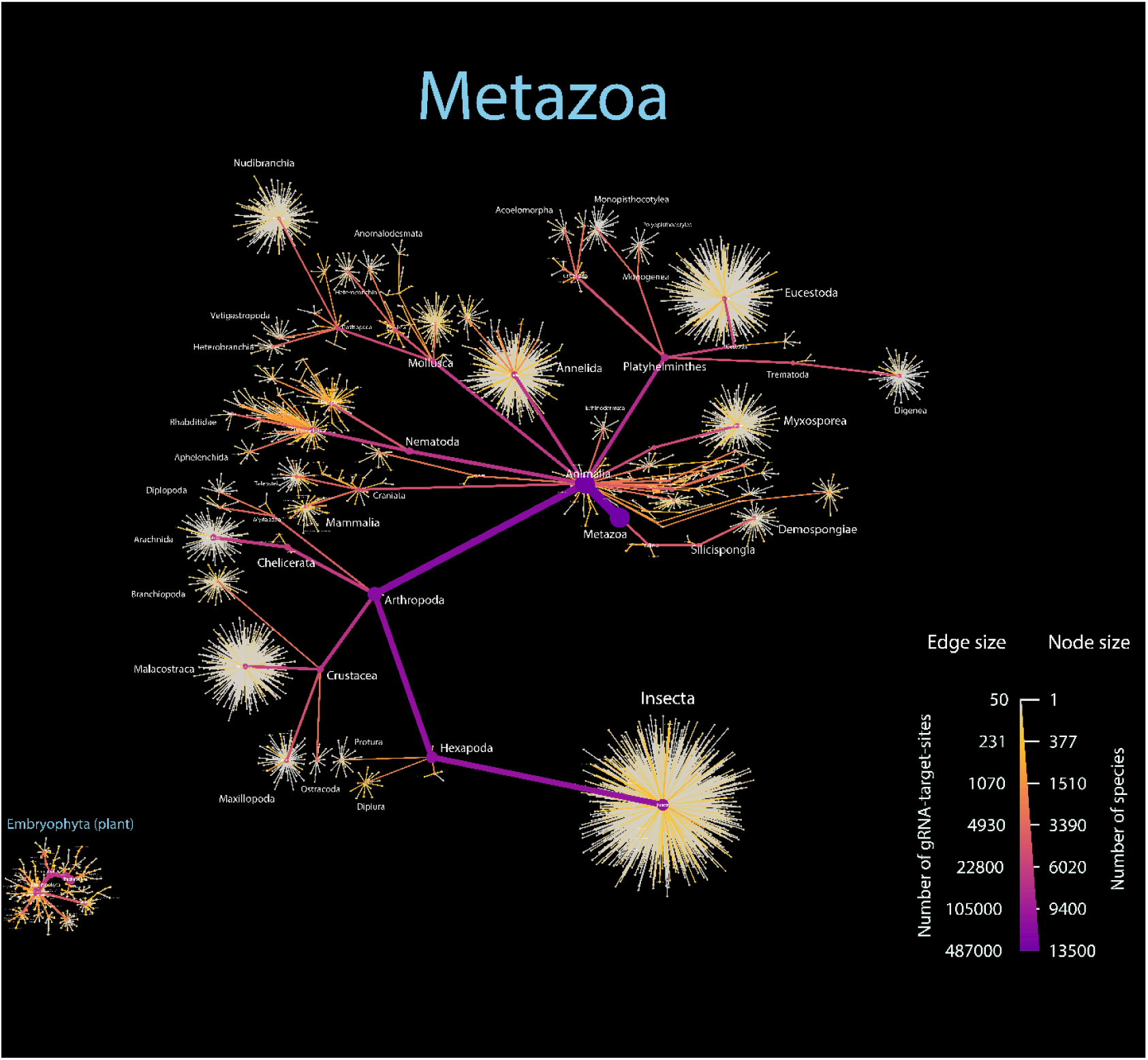
Distribution of the number of gRNA-target-sites for metazoan and plant species that are available in the SILVA SSU v119 database^38^. These gRNA-target-site oligonucleotide sequences are used for designing and synthesizing the taxon-specific and the CRISPR-Cas9 compatible sgRNA with the aim to specifically cut the 18S rRNA genes of a metazoan or plant host but not microeukaryotes (protists and fungi) with using CRISPR-Cas9. The node size indicates the number of species at each corresponding taxonomic level, while the size of the edge presents the number of gRNA-target-sites. Nodes and edges size with the highest value are coloured in purple, while the smallest size is in grey colour. Only these taxa that contain more than 50 gRNA-target-sites per taxon are showed and plotted in this image.

## Discussion

CCSAS provides a new way to obtain high-resolution taxonomic data for the eukaryotic microbiomes of plants, animals and other metazoa. By employing CRISPR-Cas9 with taxon-specific gRNAs, the background of host 18S sequences is greatly reduced or eliminated; thus, only a fraction of the sequencing effort is required to obtain high-resolution taxonomic data for the eukaryotic microbiome. Moreover, the creation of a database of gRNA-target-sites, and the primary gRNA-target-site oligonucleotide design functions of the CasOligo package, makes it easy to profile the eukaryotic microbiome of metazoa and plants. Additionally, the CasOligo package provides an oligonucleotide design function, Cas9.oligo.search2(), that can be used to design custom sgRNA for any gene for which the sequence is known, and for which there is a reference database for comparison, so that the specificity of the sgRNA can be ascertained. This includes genes encoding other regions of rRNA, such as the 16S and 23S rRNA genes, or metabolic genes (e.g. COX1). Thus, CCSAS makes it possible to study the genetic diversity of any gene in complex systems, including genes that are rare, by removing any sequence that would otherwise dominate the data. The sequence-specific removal of any amplicon has a wide range of applications, including pathogen diagnosis, and studies of symbiosis and microbiome therapy.

There are a few considerations in applying CCSAS to microbiome studies. First, there are examples of gRNA recognizing the wrong target^39-40^, which might lead Cas9 to cut some protistan and fungal sequences, or incompletely cleave host sequences. As well, incomplete removal of cut host amplicons by SPRI magnetic beads might reduce sequencing efficiency. Other more efficient size-selection methods may overcome this potential difficulty. Despite these caveats, in our hands CCSAS was far superior to other currently used approaches for obtaining high-resolution eukaryotic microbiome data.

## Methods

Methods, including statements of data availability and any associated accession codes and references, are available at the online Supplementary Information.

## Acknowledgments

V. Au, M.L. Berbee, J. Caleta, M. Edgley, G. Hu, J. Kronstad, D.G. Moerman, G. Xiang, C. Yang and Y. Zhang of the University of British Columbia, and K.M. Miller from Fisheries and Oceans, Canada, are gratefully acknowledged for providing organisms for testing (Supplementary Table 1). This work was supported by the Gordon and Betty Moore Foundation (grant: GBMF#5600) and an infrastructure award from the Canada Foundation for Innovation and British Columbia Knowledge Development Fund to C.A.S. A.C. was supported by an award from the Pacific Research Board (grant: NPRB GSRA#1335).

## Author Contributions

During a stimulating discussion to profile the eukaryotic microbiota associated with pacific oysters, C.M.D. proposed the idea of using CRISPR-Cas9 technology to reduce the background of 18S rRNA genes and conducted a pilot trial. K.X.Z. conceived, developed and implemented the CCSAS method, the CasOligo package, and the database of gRNA-target-sites. A.C. contributed to the optimization of the CCSAS method. C.A.S and A.M.C. were involved in discussions of experimental design and interpretation of the data. K.X.Z and C.A.S. wrote the manuscript with input from all authors.

## Additional information

**Supplementary Information** is available in the online version of the paper.

**Reprints and permissions information** is available at www.nature.com/reprints.

**Correspondence and requests for materials** should be addressed to C.A.S. or K.X.Z.

**Publisher’s note** Springer Nature remains neutral with regard to jurisdictional claims in published maps and institutional affiliations.

## Supplementary Information

### Supplementary Materials and Methods

#### Organisms and samples

Ten model organisms, human (*Homo sapiens*), salmon *(Salmo salar*), shrimp (*Solenocera crassicornis*), chicken (*Gallus gallus domesticus*), cow (*Bos taurus*), mouse (*Mus musculus*), fruit fly (*Drosophila melanogaster*), rock cress (*Arabidopsis thaliana*), oyster (*Crassostrea gigas*) and the nematode (*Caenorhabditis elegans*), as well as nine different species of protists and fungi were obtained from either commercial markets or various laboratories at University of British Columbia (Supplementary Table 1). As well, eight samples of seven-to 28-day old oyster spat, with sizes rangeing between 400 and 1000 μM, were obtained from a hatchery that was experiencing mortality events. The oyster spat were immediately frozen using liquid nitrogen following collection, and stored at −80°C until analysis.

#### Genomic DNA extraction

DNA from the model organisms, protists, fungi and oyster spat were extracted using the DNeasy Blood & Tissue Kit (Qiagen) following the manufacturer’s directions, and quantified using the Qubit™ DNA HS Assay Kit (Invitrogen).

To make a mock community of microeukaryotes, equal amounts of DNA from each protist and fungus (Supplementary Table 1) were pooled.

#### Design and synthesis of taxon-specific sgRNA

The specificity of CRISPR-Cas9 is determined by a 20-nt guide sequence within the sgRNA, which directs Cas9 to cut a target DNA at the 20-nt target site that is complementary to this guide sequence. Thus, the design of a taxon-specific sgRNA requires identifying a 20-nt gRNA-target-site oligonucleotide sequence in the host 18S rRNA gene, which is, absent in microeukaryotes. This taxon-specific 20-nt gRNA-target-site oligonucleotide sequence, reverse-complement to the sgRNA’s guide sequence, determines the specificity of the sgRNA and thereby the CRISPR-Cas action that is to cut 18S rDNA from the host but not from microeukaryotes. This taxon-specific 20-nt gRNA-target-site oligonucleotide sequence is used to synthesize the taxon-specific sgRNA in the laboratory using the NEB EnGen™ sgRNA Synthesis Kit.

##### Obtaining the host 18S rRNA gene sequences

Prior to the design of the sgRNA, we need to obtain the 18S rRNA gene sequence of the host organism for the identification of the gRNA-target-sites. In this study, we employed the cloning-sequencing approach as follows.

For each host, 18S rRNA gene fragments were PCR amplified using the “universal” primers TAReuk454FWD1 and TAReukREV3^26^, and sequenced to facilitate the design of gRNA-target-site oligos targeting each host. Briefly, PCR was conducted in four separate reactions run at annealing temperatures of 45, 47, 48 or 49°C, to ensure amplification of a 380-450 bp fragment from the V4 region of the 18S rRNA gene. Each 25 μL reaction mix was made with 1X PCR buffer, 4 mM MgCl_2_, 50 µg of Bovine Serum Albumin (Invitrogen), 200 mM of each dNTP (Invitrogen), 0.4 µM of each primer, 0.5 U of Q5® high fidelity polymerase (NEB) and 10 ng of genomic DNA template. As previously described^26^, the initial denaturation and activation was at 95°C for 5 min, followed by 10 cycles consisting of 95°C for 30 s, 57°C for 45 s, and 72°C for 1 min, followed by 25 cycles of denaturation at 95°C for 30 s, annealing at 45, 47, 48 or 49°C for 45 s, elongation at 72°C for 60 s, and a final elongation for 10 min at 72°C. The PCR products from the four reactions were then pooled, and the 18S amplicons purified using Agencourt SPRI magnetic beads (Beckman Coulter) at a 1:1 (vol:vol) ratio of beads:DNA to remove fragments <200bp.

These purified amplicons were then cloned into pCR2-TOPO vectors using the TOPO TA Cloning Kit (Invitrogen). Four 18S rRNA gene clones from each model organism were sent for Sanger sequencing at the NAPS Unit sequencing facility at The University of British Columbia. The obtained DNA sequences were then used to design the taxon-specific 20-nt gRNA-target-site sequence, which was be used for synthesizing the taxon-specific sgRNA that guides Cas9 to cleave the host 18S sequences, as outlined below.

##### Design of the taxon-specific gRNA-target-site oligonucleotide sequences

We developed the R package CasOligo (https://github.com/kevinzhongxu/CasOligo) for designing taxon-specific 20-nt gRNA-target-site oligonucleotide sequences, which allow sgRNA to recognize 18S sequences from specific taxa. Taxon-specific gRNA-target-site oligonucleotide sequences were designed for each model organism using the Cas9.gRNA.oligo1() function in CasOligo by providing a fasta file of the V4 region of the 18S rRNA gene from each organism that is amplified by the “universal” 18S primers, TAReuk454FWD1 and TAReukREV3^26^. The same approach can be used to design taxon-specific gRNA-target-site oligonucleotide sequences for any host organism. First, Cas9.gRNA.oligo1() searches the forward and reverse strands of the 18S rRNA gene for 20-nt gRNA-target-site oligonucleotide sequences that are compatible with Cas9 nuclease; compatibility requires that the protospacer-adjacent-motif (PAM), NGG, is immediately adjacent to the 3’ downstream region of the 20-nt target-site sequence. Each of these 20-nt gRNA-target-site sequences is potentially a target for the combined actions of sgRNA and Cas9. Next, each potential gRNA-target-site sequence is searched against the SILVA SSU database for the V4 region of 18S rRNA genes, in order to determine if the sequence is absent in protistan and fungal microeukaryotes. If so, this gRNA-target-site sequence can be used to synthesize a sgRNA that will guide Cas9 to specifically cut the host 18S rDNA. The gRNA-target-site oligonucleotide sequences designed in this study are shown in Supplementary Table 2.

##### Synthesis of sgRNA-template oligonucleotides

Once suitable taxon-specific 20-nt gRNA-target-site oligonucleotide sequences were identified, the sgRNA-template oligonucleotide sequences were obtained using the EnGen™ sgRNA Template Oligo Designer (https://nebiocalculator.neb.com/#!/sgrna), which adds a T7 promoter sequence at the 5′ end, and a 14-nt overlap sequence at the 3′ end of the 20-nt gRNA-target-site sequence. For our studies, this sgRNA-template oligonucleotide was synthesized by Integrated DNA Technologies (IDT), and diluted to 1 µM with molecular grade ultrapure water (Invitrogen).

##### Synthesis of sgRNA

The 1 µM sgRNA-template oligonucleotide was used as a DNA template to synthesize the sgRNA using the EnGen™ sgRNA Synthesis Kit, *S. pyogenes* (New England Biolabs) by following the manufacturer’s instructions. The resulting sgRNA was treated with amplification grade DNase I (Invitrogen) at room temperature for 15 min to remove any remaining DNA, and then purified using RNA Clean & Concentrator-25 Kit (Zymo Research) by following the manufacturer’s instructions. Finally, the fragment size of the sgRNA was assessed using an Agilent RNA 6000 Pico Kit (Agilent) and its concentration measured using a Qubit™ RNA HS Assay Kit (Invitrogen).

#### Validation of the design of taxon-specific sgRNA

To validate the design of gRNA for taxon-specific cleavage, we first generated 18S amplicons for each model organism and the mock community of protists and fungi. Then, these 18S amplicons were used to ascertain the effect of CRISPR-Cas9, in conjunction with taxon-specific sgRNA, on cleavage of the amplicons. The results were visualized on a gel using a Bioanalyzer (Agilent) and assessed using quantitative PCR (qPCR) as detailed below.

##### Preparation of the host 18S amplicons

For each of the ten host organisms and the mock community of protists and fungi, 18S rRNA gene fragments were obtained using PCR with the “universal” primers TAReuk454FWD1 and TAReukREV3^26^ following the conditions as detailed above. The 18S amplicons were purified using Agencourt SPRI magnetic beads (Beckman Coulter) at a 1:1 (vol:vol) ratio of beads:DNA.

##### DNA cleavage using CRISPR-Cas9

For each of the ten host organisms and the mock community of protists and fungi, the purified 18S amplicons were cut using Cas9 Nuclease, *S. pyogenes* (New England Biolabs) in the presence of a sgRNA, following the manufacturer’s directions. Briefly, the 10 µL reaction contained approximately 0.1 pmol of dsDNA, 1 pmol of sgRNA, and 1 pmol of Cas9, as well as 1x Cas9 reaction buffer to keep the molar ratio of Cas9:sgRNA:template DNA at 10:10:1. The reaction was incubated at 37°C for 4 h in a thermocycler, and then followed by 70°C for 10 min to deactivate the CRISPR-Cas9. For each sample, in parallel with the CRISPR-Cas9 treatment we also prepared the reaction without CRISPR-Cas9 treatment, in which Cas9 nuclease and sgRNA were replaced with molecular grade ultrapure water (Invitrogen). Thus, each reaction of both treatments contained the same amount of template dsDNA (18S amplicons at 0.1 pmol) and was subjected to the same incubation conditions.

##### Visualization using gel electrophoresis

The size of the 18S rDNA fragments with and without CRISPR-Cas9 treatment was visualized by gel electrophoresis using a Bioanalyser (Agilent). Prior to loading into the gel, the Cas9-cut product (5 µL out of 10 µL) were treated with 1 mg/mL (final) Proteinase K (Invitrogen) at room temperature for 15 min to digest the Cas9 nuclease. Then, 1 to 2 µL of this proteinase-K-treated product was added into a well of an Agilent High Sensitive DNA Chip in a Bioanalyzer (Agilent) to visualize and verify cutting by CRISPR-Cas9.

##### Quantitative PCR

To determine the efficiency of CRISPR-Cas9 for eliminating host-derived 18S sequences, we used quantitative PCR (qPCR) and the primer set TAReuk454FWD1/TAReukREV3^26^ (Supplementary Table 3) that targets a 380-450 bp fragment of the V4 region of the 18S rRNA gene, to assess the proportion of 18S amplicons cut by Cas9. The 10 µL qPCR reactions contained 1 X SsoFast™ EvaGreen® Supermix (Bio-Rad), 0.5 µM of each primer, and a 1 µL 1/10000 dilution of DNA template consisting of amplified products, either with or without the addition of Cas9. Thermal cycling was conducted in a CFX96 real-time PCR detection system (Bio-Rad) with the following program: 3 min denaturation at 95°C, followed by 40 cycles of denaturation at 95°C for 30 s, and annealing and extension at 49°C for 30 s. Nine, 10-fold serially diluted standards (ranging from 5 × 10^0^ to 5 × 10^9^ molecules per mL) were run in duplicate along with two no-template control reactions containing 1 µL of nuclease-free water. The amplicon standards were made from a cloned 18S rRNA gene fragment amplified from a culture of the prasinophyte microalga, *Micromonas pusilla*, using the primer set TAReuk454FWD1/TAReukREV3^26^. The amplicons were purified using a MiniElute® PCR Purification Kit (Qiagen), and quantified using a Qubit® dsDNA High Sensitivity Assay Kit (Invitrogen). The size of the amplicon was checked using gel-electrophoresis, and the qPCR melting curves were used to confirm that the fluorescence signal corresponded to a single-sized DNA fragment. The qPCR amplification efficiency was between 0.95 and 1.05 for the cloned amplicons (with r>0.98, n=9).

#### Sequencing library preparation using CRISPR-Cas Selective Amplicon Sequencing (CCSAS)

To profile host-associated eukaryotic microbiomes, we developed CRISPR-Cas Selective Amplicon Sequencing (CCSAS), which combines the use of CRISPR-Cas9 and universal 18S primers to prepare a sequencing library that is compatible with Illumina sequencing platforms. The method uses a taxon-specific sgRNA to guide Cas9 nuclease to selectively cleave 18S rRNA gene sequences from metazoa and plants, which then can be removed by size selection with SPRI beads; sequences from microeukaryotes are left intact, and can be amplified by PCR. Therefore, CCSAS allows high-resolution profiling of host-associated eukaryotic microbiomes with relatively low sequencing effort. In this study, we present CCSAS (Fig. 2); the two-step CRISPR-Cas procedure first uses Cas9 to cut the host gene encoding 18S rDNA, followed by a second cut of any host-derived 18S amplicons. Details of the method are provided below.

##### Cas9 cut on host genomic DNA

Genomic DNA of the host is cut using Cas9 Nuclease, *S. pyogenes* (NEB) following the manufacturer’s directions. Briefly, a 10-µL reaction mix containing approximately 0.1 pmol of genomic DNA, 1 pmol of sgRNA and 1 pmol of Cas9, as well as 1x Cas9 reaction buffer to keep the molar ratio of Cas9:sgRNA:template DNA at 10:10:1 is incubated at 37°C for 4 h in a thermocycler.

##### The first PCR and size selection

The Cas9-cleaved genomic DNA is used as a template in the first PCR to generate 380-450 bp amplicons from the V4 region of the 18S rDNA gene that are depleted in host sequences. To ensure representative amplification of 18S sequences from microeukaryotes, four parallel PCR reactions are run at different annealing temperatures (45, 47, 48 or 49°C), using the “universal” 18S primers TAReuk454FWD1-Nxt and TAReukREV3-Nxt (Supplementary Table 3). Compared to TAReuk454FWD1 and TAReukREV3^26^, this modified primer set contained overhang adapter sequences (Supplementary Table 3), which are compatible with Illumina indexes and sequencing adapters. These adapters allowed for a second PCR to append Illumina Nextera XT indexes to each side of the amplicons as forward and reverse primers, thus creating a dual-indexed library. This dual-indexed library preparation approach is adapted from the ref 41.

Details on the first PCR reactions are as follows. Briefly, each 25 μL reaction mix contained 1X PCR buffer, 4 mM MgCl_2_, 50 µg of Bovine Serum Albumin (Invitrogen), 200 mM of each dNTP (Invitrogen), 0.4 µM of each primer, 0.5 U of Q5® high fidelity polymerase (NEB) and 5 μL of the Cas9-cleaved genomic DNA. Because the reverse primer is 2bp shorter than the forward primer and has a lower annealing temperature, we used the two-step PCR approach of Stoeck et al.^26^, in which there is an initial ten PCR cycles at an annealing temperature where only the forward primer will bind and amplify, followed by 25 cycles at one of four lower annealing temperatures (45, 47, 48 or 49°C) where both forward and reverse primers amplify. The program has an initial denaturation and activation at 95°C for 5 min, followed by ten three-step cycles consisting of 94°C for 30 s, 57°C for 45 s, and 72°C for 1 min, followed by 25 cycles of denaturation at 94°C for 30 s, annealing at either 45, 47, 48 or 49°C for 45 s and elongation at 72°C for 60 s, with a final elongation for 10 min at 72°C. In the end, the PCR product of four reactions are pooled together. Then amplicons are size-selected and purified using magnetic Agencourt SPRI beads (Beckman Coulter) at an 0.8:1 (vol:vol) ratio of beads:DNA to remove fragment < 300bp.

##### Cas9 cut on 18S amplicons

To further remove 18S host amplicons, the size-selected amplicons described above were cut again using Cas9 Nuclease, *S. pyogenes* (NEB). Briefly, the 10 µL reaction contained approximately 0.1 pmol of DNA amplicons, 1 pmol of sgRNA, 1 pmol of Cas9, 1x Cas9 reaction buffer to keep the molar ratio of Cas9:sgRNA:template DNA at 10:10:1. The reaction was incubated at 37°C for 4 h in a thermocycler.

##### The 2^nd^ PCR and size selection

The product of the second Cas9-cut was used as the template for a second PCR (index PCR) to generate the indexed amplicons libraries. The 50-μL reaction mix of the second PCR comprised 1X PCR buffer, 4 mM MgCl_2_, 200 mM of each dNTP (Invitrogen), 5 µL of each index primer (N7XX and S5XX of Nextera® XT Index Kit), 1 U of Q5® high fidelity polymerase (NEB) and 5 µL of the product of the second Cas9 cut. The second PCR (index PCR) consisted of an initial denaturation and activation at 95°C for 3 min, followed by 29 three-step cycles consisting of 95°C for 30 s, 55°C for 30 s, and 72°C for 30 s, and a final elongation for 10 min at 72°C. The indexed amplicons generated by the second PCR were size-selected and purified using magnetic Agencourt SPRI beads (Beckman Coulter) at a ratio of 0.8:1 (vol:vol) for beads:DNA to remove fragments < 300bp.

During size selection with SPRI magnetic beads, the bead:DNA ratio depends on the size of the fragments that need to be separated. As the size of the fragments generated by cutting the ∼424-bp metazoan 18S rRNA gene sequences will vary depending on the cut site, the beads:DNA ratio of specific sgRNA may need to be optimized to remove all of the cleaved fragments. It is important to remove sequence fragments generated by amplification of the cleaved host 18S rRNA genes, as these can reduce sequencing efficiency.

#### Sequencing library preparation for amplicons generated using the universal 18S primers

Sequencing libraries for 18S amplicons generated using the “universal” 18S primers and without cutting using CRISPR-Cas9 were prepared using protocols adapted from Illumina^41^. Briefly, two successive runs of PCR were performed as follows: For the first PCR, 29 cycles of amplification using the modified primers TAReuk454FWD1-Nxt and TAReukREV3-Nxt (Supplementary Table 3) were used to generate 380 to 450 bp amplicons of the V4 region of the 18S rRNA genes. The reaction conditions for the first PCR were as detailed above for the first CCSAS PCR, except that there was about 5 ng of genomic DNA in the sample. The amplicons were purified using magnetic Agencourt SPRI beads (Beckman Coulter) at a ratio of 1:1 (vol:vol) for beads:DNA to remove fragments < 200bp.

Five µL of the purified amplicons from the first PCR were used as templates for the second PCR (index PCR). The PCR reactions and conditions for the second PCR were as detailed above for the second CCSAS PCR. The amplicon libraries generated were purified using magnetic Agencourt SPRI beads (Beckman Coulter) at a ratio of 1:1 (vol:vol) for beads:DNA to remove fragments < 200bp.

#### Sequencing library preparation for amplicons generated using the non-metazoan primers

Preparation of the sequencing library for the 18S amplicons obtained using non-metazoan primers^27^ was similar to that described above, except that the first PCR used the primers 18s-EUK581-F-Nxt and 18s-EUK1134-R-Nxt (Supplementary Table 3) to amplify a 300-600 bp region of 18S rRNA genes that is specific to microeukaryotes but absent in metazoa^27^. The 25 μL reaction mix comprised 1X PCR buffer, 4 mM MgCl_2_, 50 µg of Bovine Serum Albumin (Invitrogen), 200 mM of each dNTP (Invitrogen), 0.4 µM of each primer, 0.5 U of Q5® high-fidelity polymerase (NEB) and approximately 5 ng of genomic DNA. There initial denaturation of 2 min at 98°C was followed by 29 cycles of denaturation at 98°C for 30 s, annealing at 51.5°C for 30 s and elongation at 72°C for 60 s, and a final elongation for 10 min at 72°C. The first PCR product was purified using magnetic Agencourt SPRI beads (Beckman Coulter) at a ratio of 1:1 (vol:vol) for beads:DNA to remove fragments less < 200bp (e.g. dimers). The amplicon libraries were completed as described above, with a second PCR to add a Nextera® XT index to each 3’ and 5’ end of the amplicons.

#### Next-generation sequencing and data analysis

The DNA concentrations of the 18S amplicon sequencing libraries that were prepared using “universal” 18S primers, non-metazoan primers, or the CCSAS method were measured using the Qubit® dsDNA High Sensibility Assay Kit (Invitrogen). The fragment size for each type of library was determined using an Agilent bioanalyzer with the High Sensitive DNA Chip (Agilent). Equimolar amounts of these barcoded and purified amplicon sequencing libraries were pooled and sent for sequencing at the BRC Sequencing Core at the University of British Columbia using MiSeq Illumina 2 × 300bp chemistry.

Sequences were processed and analyzed using QIIME version 1.9^42^. Briefly, sequences were de-multiplexed by their forward and reverse indexes, and the paired-end reads merged using PEAR version 1.10.4^43^. Then, sequences from different samples were pooled, and Uclust^44^ was used for OTU picking with 99% nucleotide sequence similarity. Taxonomy was assigned for representative OTU sequences using the Uclust consensus taxonomy assigner and the SILVA SSU database^38^ (*version v132, released on 13 December 2017*) at a 90% confidence cutoff. The samples were normalized by analyzing the relative abundance for each OTU or taxon as the proportion of all sequences (tags) within a sample. The data is visualized using ggplot2^45^ and metacoder^46^.

## Data Availability

All next-generation sequencing data generated in this study have been deposited in the NCBI Sequence Read Archive (SRA) under BioProject (PRJNA625176). The Sanger cloning-sequencing data were deposited in GenBank under the accession numbers MT328571 to MT328580 for 18S rRNA gene sequences of ten model organisms. The authors declare that all other data supporting the findings of this study are available within the paper and/or the associated supplementary files. All related scripts, functions and algorithms for designing gRNA-target-site oligonucleotide sequences were included in the custom R package: CasOligo (https://github.com/kevinzhongxu/CasOligo). The gRNA-target-sites database was included in the CasOligo package as well.

---

**Supplementary Figure 1.**
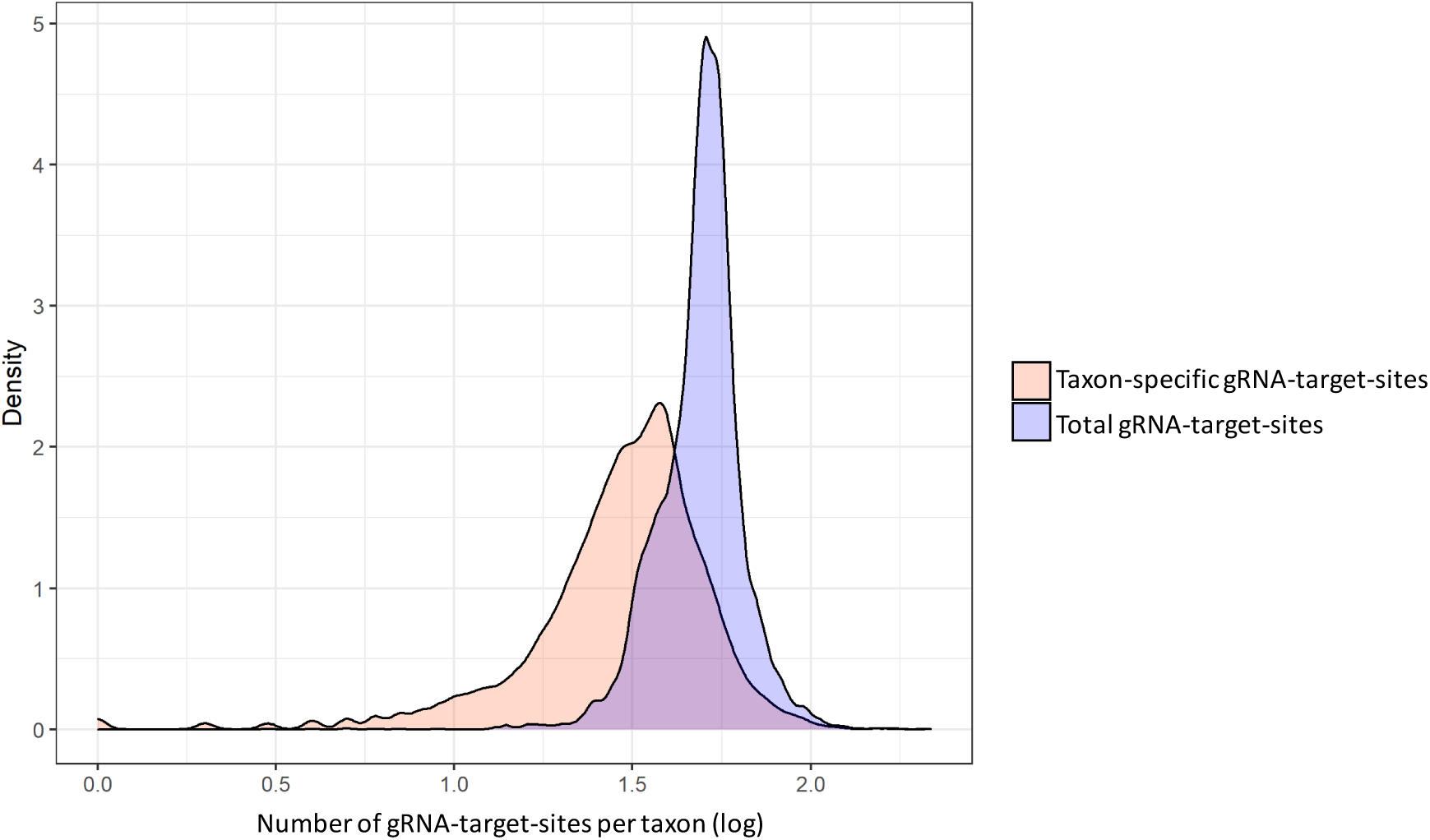
Distribution of the number of gRNA-target-sites of each metazoan and plant species from the SILVA SSU v119 database^38^. These gRNA-target-site oligonucleotide sequences were identified, using the Cas9.gRNA.oligo1() algorithm, from the V4 region of the 18S rRNA gene that is flanked by the 18S “universal” primer set TAReuk454FWD1 / TAReukREV3^26^, and are used for designing and synthesizing the CRISPR-Cas9-compatible sgRNA. The taxon-specific gRNA-target-sites allows the design of the sgRNA to taxon-specifically cut the 18S rRNA gene sequence of a metazoan or plant host but not microeukaryotes (protists and fungi) using CRISPR-Cas9.

**Supplementary Figure 2.**
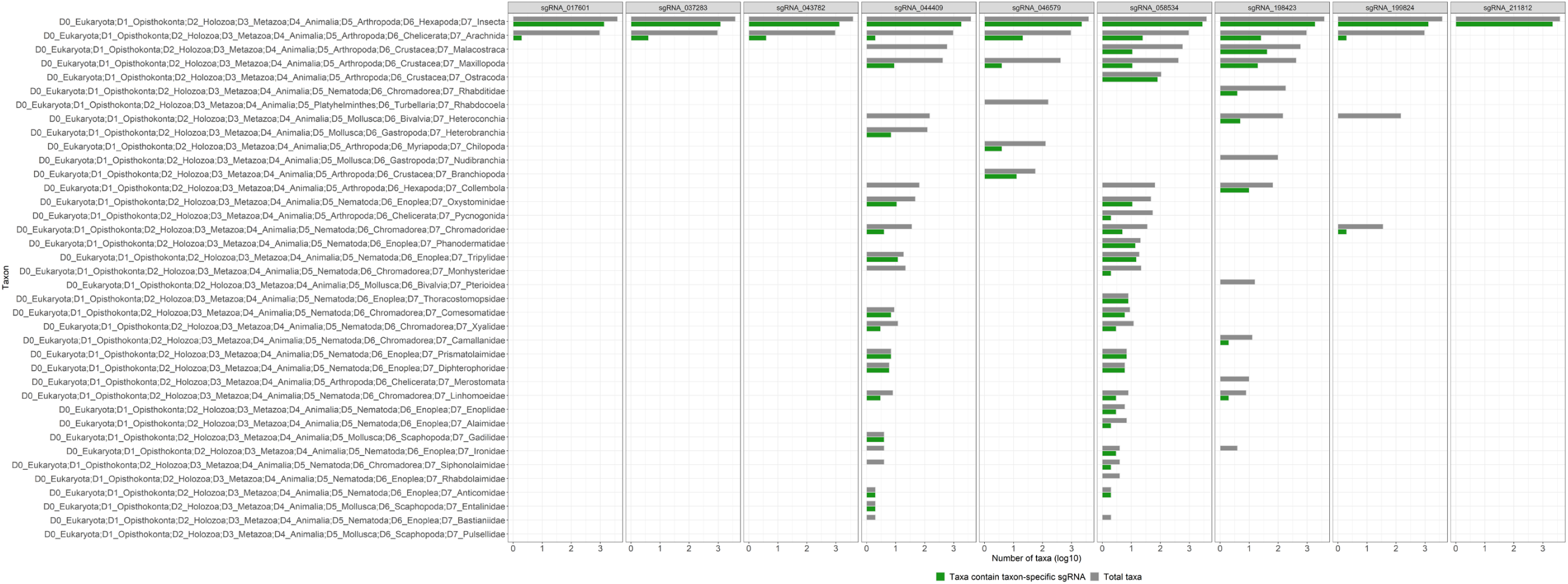
Summary of the number of eukaryotic species at each D7 taxonomic level that the sgRNA can cut at the V4 region of the 18S rRNA genes that are flanked by the 18S “universal” primer set TAReuk454FWD1 / TAReukREV3^26^. These nine sgRNAs, which are among 205242 unique taxon-specific sgRNA designed from the SILVA SSU database (*version 119*)^38^ using CasOligo, are selected to show that some sgRNAs can target more than 1000 species and broad taxonomic groups based on an *in-silico* analysis (*i*.*e*. 100% match to the 18S rRNA gene sequences of the metazoan host at the gRNA-target-site, but no match for protists and fungi). Taxon names on the left side of the panel are shown as SILVA taxonomic hierarch with levels range from D0 (kingdom) to D7. The D7 taxonomic level comprises eukaryotic classes and families.

**Supplementary Table 1.** List of organisms used in this study.

| Organisms | Classification | Representative groups | Applications | Source | NCBI accession no. |
| --- | --- | --- | --- | --- | --- |
| *Cryptococcus neoformans* variety grubii | Fungi, Basidiomycota | Fungi | Mock community of protists-fungi | J. Kronstad, UBC | - |
| *Ustilago maydis* strain 002 WT | Fungi, Basidiomycota | Fungi | Mock community of protists-fungi | J. Kronstad, UBC | - |
| *Aspergillus citrisporus* strain E1 | Fungi, Ascomycete | Fungi | Mock community of protists-fungi | M.L. Berbee, UBC | - |
| *Rhizopus* sp. strain AFTDL | Fungi, Zygomycete | Fungi | Mock community of protists-fungi | M.L. Berbee, UBC | - |
| *Batrachochytrium dendioatriodio* strain JEL 423 | Fungi, Chytride | Fungi | Mock community of protists-fungi | M.L. Berbee, UBC | - |
| *Coemunsia* sp. | Fungi, Zygomycete | Fungi | Mock community of protists-fungi | M.L. Berbee, UBC | - |
| *Creolimax fragrantissima* strain Pdd4 | Protist, Ichthyosporaea | Protists | Mock community of protists-fungi | M.L. Berbee, UBC | - |
| *Micromonas pusilla* | Protist, Mamiellaceae | Protists | Mock community of protists-fungi | This lab | - |
| *Tetrahymena* sp. | Protist, Tetrahymenidae | Protists | Mock community of protists-fungi | This lab | - |
| *Caenorhabditis elegans* | Animalia, Nematoda, Caenorhabditis | Worm | Model organisms | D.G. Moerman, UBC | MT328580 |
| *Arabidopsis thaliana* | Plantae, Angiosperms, Arabidopsis | Plant | Model organisms | Y. Zhang, UBC | MT328579 |
| *Homo sapiens* (Human) | Animalia, Chordata, Mammals | Mammals | Model organisms | The skin tissue of my lip | MT328572 |
| *Mus musculus* (Mouse) | Animalia, Chordata, Mammals | Mammals | Model organisms | J. Caleta, UBC | MT328573 |
| *Bos taurus* (Cow) | Animalia, Chordata, Mammals | Mammals | Model organisms | Commercial market | MT328574 |
| *Gallus gallus* (Chicken) | Animalia, Chordata, Aves | Aves | Model organisms | Commercial market | MT328575 |
| *Salmo salar* (Salmon) | Animalia, Chordata, Actinopterygii | Fish | Model organisms | Commercial market | MT328576 |
| *Solenocera crassicornis* (Shrimp) | Animalia, Arthropoda, Malacostraca | Malacostraca | Model organisms | Commercial market | MT328577 |
| *Crassostrea gigas* (Oyster) | Animalia, Mollusca, Bivalvia | Bivalvia | Model organisms | K.M. Miller, Fisheries & Oceans, Canada | MT328571 |
| *Drosophila melanogaster* | Animalia, Arthropoda, Insecta | Insect | Model organisms | Wild nature | MT328578 |

**Supplementary Table 2.**
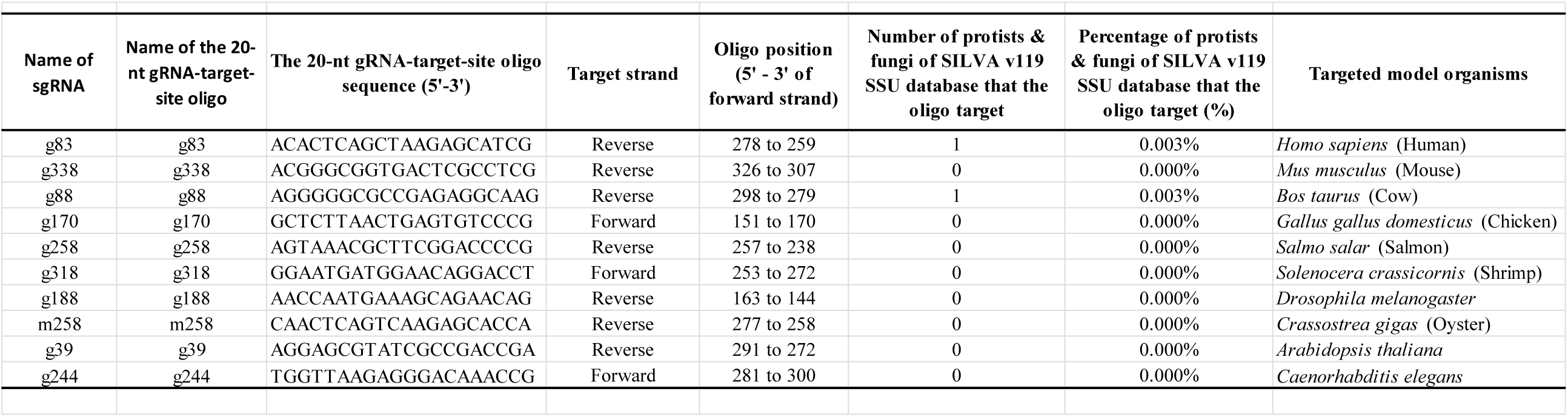
List of the 20-nt sgRNA-target-site oligonucleotide sequences designed for cutting V4 region of 18S rRNA genes of ten model organisms using CRISPR-Cas9.

**Supplementary Table 3.**
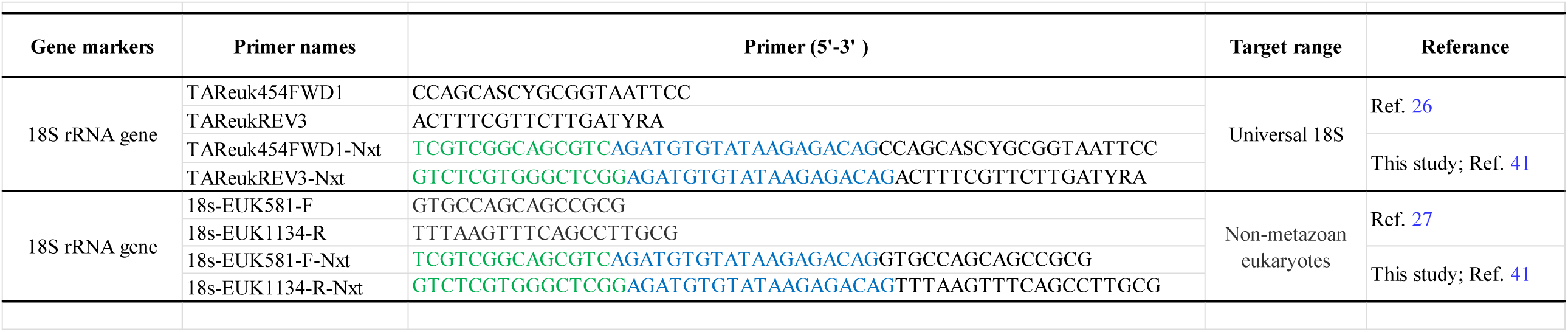
List of primers used in this study.

